# Phosphorylation of the α2 glycine receptor induces an extracellular conformational change and slows the rise and decay rates of glycinergic synaptic currents

**DOI:** 10.1101/434357

**Authors:** Sharifun Islam, Xiumin Chen, Argel Estrada-Mondragon, Joseph. W. Lynch

## Abstract

The α2 glycine receptor (GlyR) is a pentameric ligand-gated anion channel that plays a key role in cortical interneuron migration and in the differentiation of cortical progenitor cells into functional neurons. It also mediates tonic inhibitory chloride currents in adult forebrain neurons. Disruption of α2 GlyR gene expression or receptor function results in the aberrant functioning of neuronal circuits which contributes to the pathophysiology of schizophrenia, autism and epilepsy. This implicates the α2 GlyR as a possible therapeutic target for a range of neurological disorders. However, despite its therapeutic potential, little is known about the mechanisms by which α2 GlyRs are functionally modulated. To address this, we investigated whether the α2 GlyR is modulated by phosphorylation at a serine residue (S341) within the same PKA consensus sequence (R-E-S-R) that houses the α3 GlyR S346 residue that is known to be phosphorylated by PKA. Resolving this question might uncover a novel means of physiologically, pathologically or therapeutically modulating α2 GlyRs. We show using voltage-clamp fluorometry that forskolin-induced phosphorylation of S341 induces a conformational change in the glycine binding site. We also employed glycinergic artificial synapses to demonstrate that the S341E phospho-mimetic mutation slows the rise and decay rates of α2-mediated glycinergic inhibitory postsynaptic currents. These results suggest that PKA phosphorylation alters the structural and functional properties of the α2 GlyR. This information may help to identify new mechanisms by which α2 GlyRs may be pathologically modified or therapeutically targeted for the treatment of neurological disorders.

## Introduction

Ligand-gated ion channels (LGICs) allow cells to respond rapidly to changes in their external environment. Members of the pentameric ligand-gated ion channel (pLGIC) receptor family are composed of five protein subunits that form a pentameric arrangement around a central ion conducting pore. The major vertebrate members of the family include the cation permeable nicotinic acetylcoline receptor (nAChR) and serotonin type-3 receptor (5HT3R), and the anion permeable GABA type-A (GABAAR) and glycine receptor (GlyR). Each subunit consists of large N-terminal ligand-binding domain that incorporates orthosteric ligand-binding sites at the interface of adjacent domains. The ligand-binding domain is coupled to a transmembrane domain that comprises four transmembrane α-helices, termed M1-M4, that each span the full thickness of the cell membrane. Each subunit contributes an M2 domain to the lining of the axial water-filled pore. The M1, M3 and M4 domains are arranged concentrically around the M2 domain bundle and thereby help to isolate the water-filled pore from the non-polar lipid membrane. M1, M2 and M3 are connected by short intracellular and extracellular loops, respectively, whereas M3 and M4 are connected by a large intracellular domain that varies considerably in both length and amino acid sequence among different family members. The M3-M4 domain contains phosphorylation sites and binding sites for synaptic clustering and trafficking proteins.

GlyRs mediate fast inhibitory synaptic transmission in neurons of the spinal cord, brain stem, caudal brain and retina [1–3]. A total of five GlyR subunits are known: α1-α4 and β. However, the human α4 subunit is a pseudogene due to a premature stop codon at the intracellular end of M4 that prevents the expression of full length transcripts [4]. All other α subunits function as homomeric receptors when recombinantly expressed, but the β subunit is functional only when incorporated with α subunits into αβ heteromeric receptors. Synaptic GlyRs comprise heteromeric αβ GlyRs, with the β subunit required for synaptic localization due to its ability to bind to the subsynaptic clustering protein, gephyrin [5–7].

The α1β GlyR is the dominant isoform in the adult central nervous system. It is best known for mediating glycinergic inhibitory transmission in adult spinal cord motor reflex arcs [2]. Consistent with this role, hereditary mutations to either α1 or β genes result in a severe motor disorder termed hyperekplexia (or startle disease) that is characterised by an exaggerated reflex startle response to unexpected stimuli, which is often accompanied by temporary but complete muscular rigidity [8].

The α3β GlyR mediates inhibitory neurotransmission in spinal nociceptive neurons in superficial laminae of the spinal cord dorsal horn [9–11]. Chronic inflammatory pain sensitization is caused by a prostaglandin E_2_ (PGE_2_)-mediated activation of protein kinase A (PKA), which in turn phosphorylates α3 GlyRs at S346, leading to a diminution of glycinergic inhibitory postsynaptic current (IPSC) magnitude [10]. This removal of inhibitory drive results in the disinhibition of spinal nociceptive sensory neurons which in turn induces chronic inflammatory pain sensitization. Because of their role in restraining nociceptive neuronal firing and their sparse distribution outside the spinal cord dorsal horn, α3 GlyRs have emerged as promising therapeutic targets for chronic pain, and indeed, agents that restore (i.e., potentiate) α3 GlyR function have been shown to exhibit analgesic efficacy in animal models of chronic inflammatory pain [11–17].

GlyR α2 transcripts predominate in the neonatal and embryonic central nervous system, and replaced postnatally by those of α1 and, to a lesser extent, α3 GlyRs, by adulthood [18]. Embryonic cortical interneurons express α2 GlyRs at high levels during their initial migratory phase, knockout of these receptors results in neuronal migration defects [19]. Genetic ablation of the α2 GlyR gene also impairs the production of mature neurons from cortical progenitor cells which reduces the number of excitatory projection neurons in deep and upper layers of the cerebral cortex, thereby resulting in reduced brain size [20]. The α2 GlyR also mediates tonic inhibitory chloride currents in adult forebrain neurons [21], and disruptions to its function would alter the excitability of forebrain neuronal circuits. All these effects work in concert to disrupt neuronal circuit development and function [22] which in turn contribute to the pathophysiology of schizophrenia, autism and epilepsy [23]. Consistent with this, hereditary mutations in the human α2 GlyR gene are associated with autism spectrum disorder [24–27]. Together, this information implicates the α2 GlyR as a potential therapeutic target for a range of neurodevelopmental disorders. To date, however, there has been little attempt to identify unique features of druggable binding sites in this receptor that could be exploited in designing drugs specific for α2 over the highly homologous α1 and α3 GlyR isoforms.

Moreover, despite their importance in neurodevelopment, little is known about the physiological or pathological mechanisms of α2 GlyR modulation. For example, it is as yet unclear whether the α2 GlyR is modulated by phosphorylation. It is known, however, that the α2 GlyR incorporates a serine residue (S341) within the same PKA consensus sequence (R-E-S-R) that houses the α3 GlyR S346 residue that is known to be phosphorylated by PKA [10, 28]. Resolving this question might uncover a possible alternate means of physiologically, pathologically or therapeutically modulating α2 GlyRs.

The aim of the present study was to determine whether α2 GlyRs are phosphorylated by PKA, and if so, whether this induced a conformational change in the glycine binding site, as has previously been shown for the homologous phosphorylation site in the α3 GlyR [28]. To achieve this, we employed voltage-clamp fluorometry (VCF) [29] to directly monitor phosphorylation-mediated conformational changes in the extracellular glycine binding site. We also employed glycinergic artificial synapses [30, 31] to determine whether phosphorylation may alter the functional properties of the inhibitory postsynaptic currents (IPSCs) mediated by α2 GlyRs.

## Materials and Methods

### Chemicals

Methanethiosulfonate-rhodamine (MTSR) and 2-((5(6)-tetramethylrhodamine)carboxylamino)ethyl methanethiosulfonate (MTS-TAMRA) were obtained from Toronto Research Chemicals. Glycine, β-alanine, taurine, strychnine, forskolin, were all obtained from Sigma. Glycine, β-alanine, taurine and strychnine were dissolved in water. All other drugs were prepared as 20-100 mM stocks in dimethylsulfoxide and kept frozen at −20 °C. From these stocks, solutions for experiments were prepared on the day of recording.

### Molecular biology

For *Xenopus* oocyte recordings, the plasmid DNA encoding the human α1 and α2 GlyRs were subcloned into pGEMHE, a plasmid vector optimized for *Xenopus* oocyte expression. Site directed mutagenesis was performed using the QuikChange mutagenesis kit (Stratagene). Successful incorporation of the mutations was confirmed through automated sequencing of the entire cDNA coding region. Chimera-1 was constructed using a multiple-template-based sequential PCR protocol as previously described [32]. The join site between the α1 and α2 sequences used to create chimera-1 was located between α1 Y223/L224 and α2 Y223/L224 at the N terminal end of M1. Chimera-1 also included the α1-N203C mutation.

Ten micrograms of each cDNA was linearized by NheI or PstI and then purified by PCR-purification kit (Qiagen). The capped RNAs were transcribed from cDNA using the Ambion T7 mMessage mMachine kit, purified by RNAMinikit (Qiagen) eluted with DNA/RNAase free water and diluted to 200 ng/ μl for oocyte injection. For HEK293 cell transfections, we employed plasmid DNA encoding the human α2 GlyR (in pCIS2), the mouse neuroligin 2A splice variant (in pNice) and empty pEGFP plasmid. Site-directed mutagenesis was performed using the QuikChange mutagenesis kit, and the successful incorporation of mutation was confirmed by DNA sequencing.

### Oocyte preparation, injection and labeling

All *Xenopus laevis* handling procedures were approved by the University of Queensland Animal Ethics Committee (approval numbers: QBI/059/13/ARC/NHMRC and QBI/AIBN/087/16/NHMRC/ARC). Female *Xenopus* frogs (Xenopus Express) were anaesthetized with 5 mM MS-222 (Sigma Aldrich) and stage VI oocytes were removed from ovaries and washed thoroughly in OR-2 (82.5 mM NaCl, 2 mM KCl, 1 mM MgCl_2_, 5 mM HEPES, pH 7.4). The oocytes were then incubated in collagenase (Sigma Aldrich) in OR-2 for 2 hr at room temperature, rinsed and stored in OR-2 at 18 °C.

All oocytes were injected with 10 ng of mRNA into the cytosol. To achieve the high levels of expression required for the detection of the fluorescent signal over the background (due to oocyte autofluorescence and non-specific binding of the dye) the oocytes were incubated at 18 °C for 3-10 days after injection. The incubation solution contained 96 mM NaCl, 2 mM KCl, 1 mM MgCl_2_, 1.8 mM CaCl_2_, 5 mM HEPES, 0.6 mM theophylline, 2.5 mM pyruvic acid, 50 μg/ml gentamycin (Cambrex Corporation) and 5 % horse serum (Hycell), at pH 7.4. On the day of recording, the oocytes were transferred into ND96 (96 mM NaCl, 2 mM KCl, 1 mM MgCl_2_, 1.8 mM CaCl_2_, 5 mM HEPES, pH 7.4) and stored on ice. To label with either MTSR or MTS-TAMRA, oocytes were transferred into the labeling solution containing 10 μM of either compound in ND96 for 25 s. The oocytes were then washed and stored in ND96 for up to 6 hr before recording. All labeling steps were performed on ice.

### VCF

We employed an inverted microscope (Ti-S, Nikon Instruments) equipped with a DAPI filter set (49000, Chroma Technology) and a CFI Fluor 40X water immersion objective (N.A. 0.80, MRF07420, Nikon Instruments). A Lambda LS 175 W xenon arc lamp served as a light source and was coupled to the microscope via a liquid light guide (Sutter Instruments). Fluorescence was detected using a H7360-03 photomultiplier (Hamamatsu Photonics) coupled to a PMT400R photomultiplier subsystem (Ionoptix). Oocytes were placed securely in the bath and were continually perfused with ND96 solution. Agonists were dissolved in the same solution and were applied to the oocytes via a gravity-fed perfusion system. Currents were recorded via two-electrode voltage clamp using 0.2-2 MΩ resistance glass pipettes filled with 3 M KCl. Oocytes were voltage-clamped at −40 mV and currents were recorded using a Gene Clamp 500B amplifier (Molecular Devices). ΔI and ΔF signals were digitized at 2 kHz via a Digidata 1322A interface and pCLAMP 9.2 software (Molecular Devices).

### Primary culture of spinal neurons

Spinal neurons were prepared using methods as previously described [30]. Briefly, E15 timed-pregnant rats were euthanized via CO_2_ inhalation in accordance with procedures approved by the University of Queensland Animal Ethics Committee (QBI/142/16/NHMRC/ARC). The spinal cords were rapidly removed, triturated and plated onto poly-D-lysine-coated coverslips in a 4-well plate at a density of 8-10×10^4^ cells/well, and cultured for 3-4 wk until spontaneous inhibitory postsynaptic currents (IPSCs) could be detected. The cells were initially cultured in Dulbecco’s Modified Eagles Medium (DMEM) supplemented with 10% foetal bovine serum (DMEM-FBS). After 24 h the entire DMEM-FBS medium was replaced with Neurobasal medium including 2% B27 and 1% GlutaMAX supplements. A second (and final) feed 1 wk later replaced half of this medium with fresh Neurobasal medium. Neurons were used in co-culture experiments between 1-4 wk later.

### HEK293 cell culture, transfection and artificial synapse formation

HEK293 cells were cultured in DMEM-FBS until ~ 90% confluent. One day prior to transfection, they were trypsinized and plated onto glass coverslips in 35 mm culture dishes at a density of 5×10^3^ cells/dish. For artificial synapses, we transfected plasmid DNA encoding the α1 or α2 GlyR, EGFP and neuroligin 2A in equal ratios. Transfection was performed via a Ca^2+^ phosphate-DNA co-precipitation method for 15-20 h in a 3% CO_2_ incubator and terminated by washing cells twice with divalent cation-free phosphate buffered saline. For artificial synapse experiments, transfected cells were trypsinized the next day, centrifuged and re-suspended in Neurobasal medium (including 2% B27 and 1% GlutaMAX supplements) then seeded onto the neurons. One 35 mm dish of HEK293 cells was typically sufficient to seed four coverslips of neurons. Once seeded with HEK293 cells, the co-cultures were returned to the incubator overnight to allow artificial synapses to form between neurons and transfected HEK293 cells. Cells were used for patch-clamp recording over the following 1-3 d.

### Patch clamp electrophysiology

Cells were viewed using an inverted microscope and currents were recorded by whole-cell patch-clamp recording. Cells were perfused by an extracellular solution containing (in mM): 140 KCl, 2 CaCl_2_, 1 MgCl_2_, 10 HEPES/ NaOH and 10 glucose (pH 7.4 adjusted with NaOH). Patch pipettes had a tip resistance of 1-4 MΩ when filled with the standard intracellular solution consisting of (mM): 145 CsCl, 2 CaCl_2_, 2 MgCl_2_, 10 HEPES and 10 EGTA (pH 7.4 adjusted with CsOH). In experiments where forskolin was applied to HEK293 cells in an attempt to phosphorylate α2 GlyRs, the intracellular solution consisted of (mM): 145 CsCl, 2 CaCl_2_, 2 Mg-ATP, 0.2 Na-GTP, 10 HEPES and 10 EGTA (pH 7.4 adjusted with CsOH).

For isolated HEK293 cell whole-cell recordings, cells were voltage clamped at −40 mV and membrane currents were recorded using Multiclamp 700B, a Digidata 1440A and pClamp 10 software (Molecular Devices). Currents were filtered at 1 kHz and digitized at 2 KHz. Solutions were applied to cells via gravity forced perfusion and parallel microtubules and manual control of this system was achieved via a micromanipulator with a solution exchange time < 250 ms. For artificial synapse recordings, cells were voltage clamped at −70 mV and membrane currents were recorded using Multiclamp 700B, a Digidata 1440A and pClamp 10 software (Molecular Devices). Currents were filtered at 4 kHz and digitized at 10 KHz. Experiments were conducted at room temperature (19-22 °C). Recordings with series resistance above 20 MΩ were discarded. Capacitance of the HEK293 cells was typically 20 pF, resulting in a typical corner frequency of 398 Hz. Because this was satisfactory for our experiments, series resistance compensation was not applied. Sliding templates in Axograph X were used to identify well-isolated IPSCs. 10-90% rise times were recorded and mono-exponential simplex fits applied to the decay period of individual events, which were then averaged for each cell.

### Data analysis

EC_50_ and n_H_ values for glycine-induced activation of current (ΔI) and fluorescence (ΔF) signals were obtained using the empirical Hill equation, fitted with a non-linear least squares algorithm (SigmaPlot 12.0, Systat Software). All results are expressed as mean ± SEM. Data sets were first tested for normal distribution prior to using ANOVA tests to determine statistical significance, with p < 0.05 as the significant threshold (SigmaPlot 12.0).

## Results

### VCF experiments reveal extracellular structural differences between α1 and α2 GlyRs

VCF involves introducing a cysteine into a location of interest in an otherwise cysteine-free protein [29]. The cysteine is then reacted with an thiol-reactive fluorophore, typically containing a methanethiosulfonate (MTS) or maleimide moiety. A key requirement is that the spectral emission properties of the fluorophore varies according to the hydrophobicity of their environment. This means that protein conformational changes occurring in the vicinity of the fluorophore can be reported in real time as fluorescence changes. Here we employed VCF to compare glycine-induced conformational changes in the N-terminal domains of α1 and α2 GlyRs in the absence and presence of phosphorylation. For this purpose we selected two sites (N203C and R271C using α1 GlyR numbering), both of which have been successfully labelled by rhodamine derivatives in previous VCF studies on α1 and α3 GlyRs [28, 33, 34]. Fig 1A shows a structural model of an α1 GlyR subunit displaying the location of both sites. The α1 GlyR N203 residue lies at the tip of the loop C binding domain which closes around the bound glycine molecule to initiate channel activation [35]. The corresponding α2 GlyR residue is N210. We also investigated the α1 GlyR R271 residue (which corresponds to R278 in the α2 GlyR) as it lies distant from the glycine binding site in a region critically involved in the channel gating mechanism [35, 36] and thus reports conformational changes associated with channel gating but not binding.

**Fig 1.**
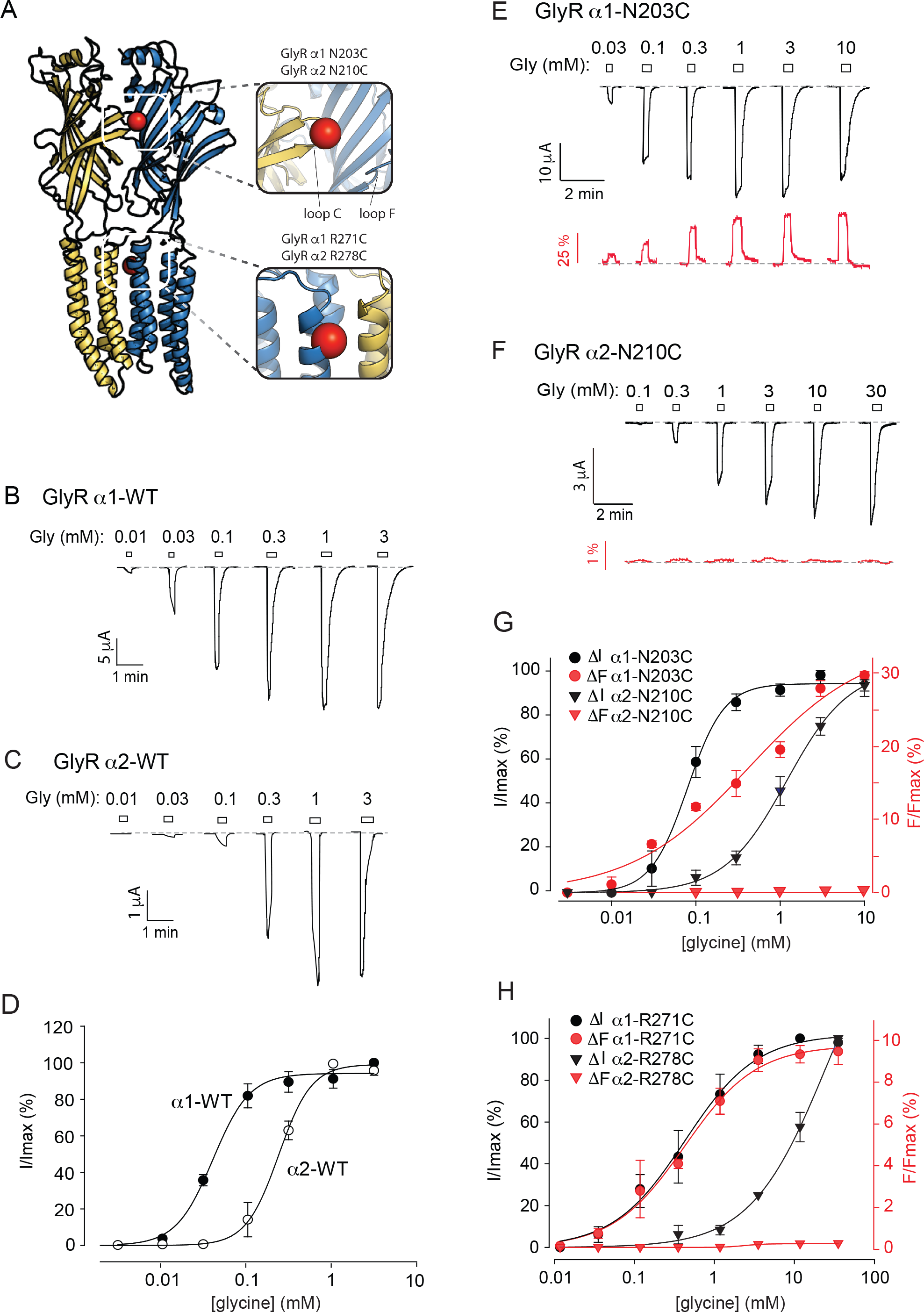
Comparison of ΔI and ΔF activation properties of WT and indicated mutant α1 and α2 GlyRs. In this and subsequent figures, ΔI and ΔF recordings are shown in black and red, respectively. **A.** Structural model of an α1 GlyR subunit showing the locations of N203 (at the tip of C domain) and R271 (at the top of M2 domain). The corresponding α2 GlyR residue numbering is also shown. **B.** Example of a α1-WT GlyR glycine ΔI dose-response relationship. **C.** Example of a α2-WT GlyR glycine ΔI dose-response relationship. **D.** Averaged ΔI and ΔF dose-response relations for the α1-WT and α2-WT GlyRs. Mean parameters of best fit to individual dose-response relations are summarized in Table 1. **E, F.** Examples of glycine EI and ΔF dose-response relationships recorded from oocytes that expressed MTS-TAMRA-labeled α1-N203C and α2-N210C GlyRs, respectively. **G.** Averaged ΔI and ΔF dose-response relations for MTS-TAMRA-labeled α1-N203C and α2-N210C GlyRs. Mean parameters of best fit to individual dose-response relations are summarized in Table 1. **H.** Averaged ΔI and ΔF dose-response relations for MTSR-labeled α1-R271C and α2-R278C GlyRs. Mean parameters of best fit to individual dose-response relations are summarized in Table 1.

Examples of glycine-activated current (ΔI) dose-response relationships for wild type (WT) α1 and α2 GlyRs are shown in Fig 1B and C with mean parameters of best fit to individual dose-response relationships summarized in Fig. 1D and Table 1. The α2-WT GlyR exhibits a dramatically higher EC_50_ value than the α1-WT GlyR (230 ± 1 11 versus 39 ± 1 μM, n = 5 each, p < 0.001), consistent with previous studies [37] [38].

**Table 1.**
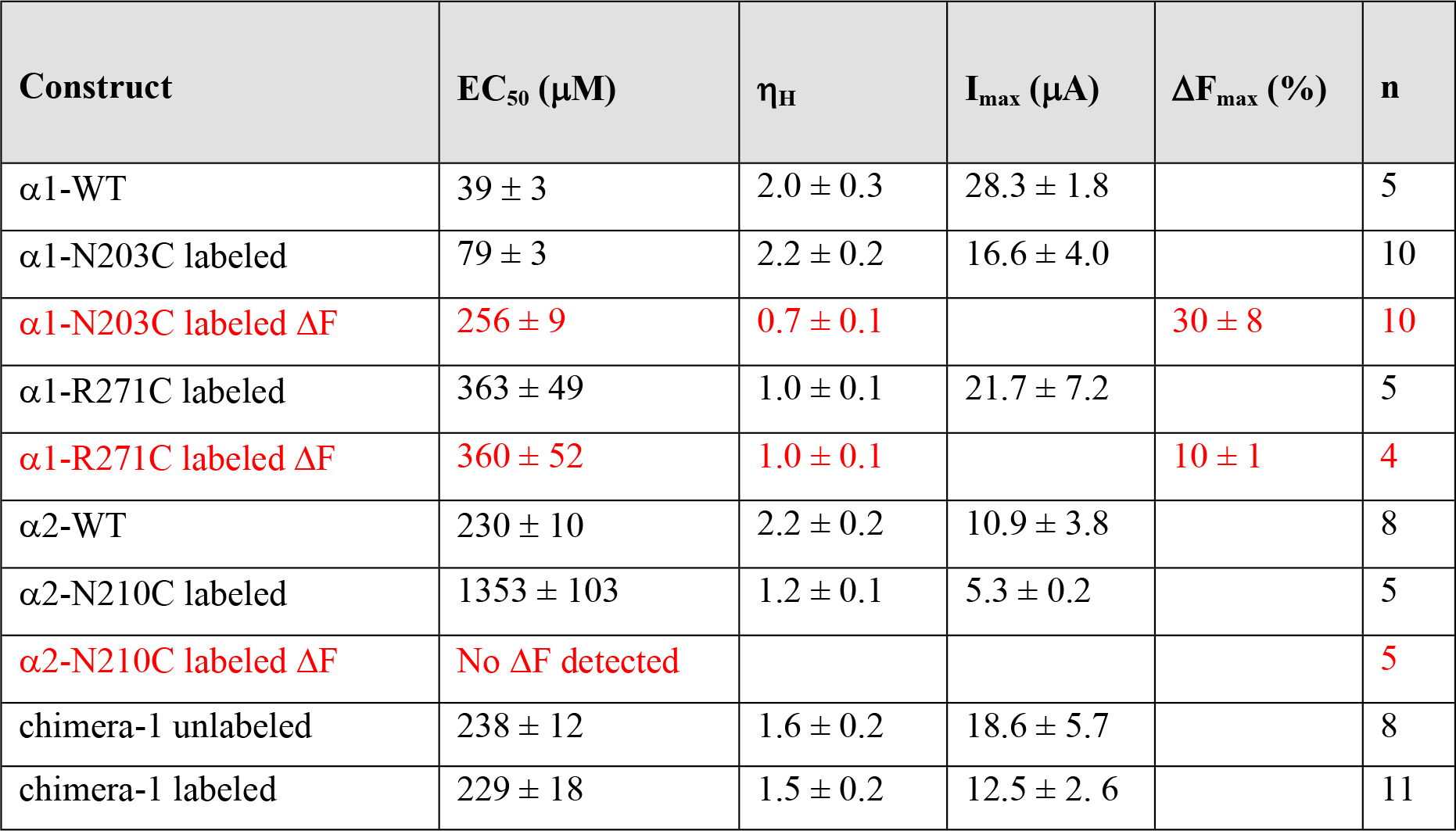
Glycine-dependent activation properties of the GlyR constructs employed in VCF experiments. Black and red text denote eI and ΔF responses, respectively.

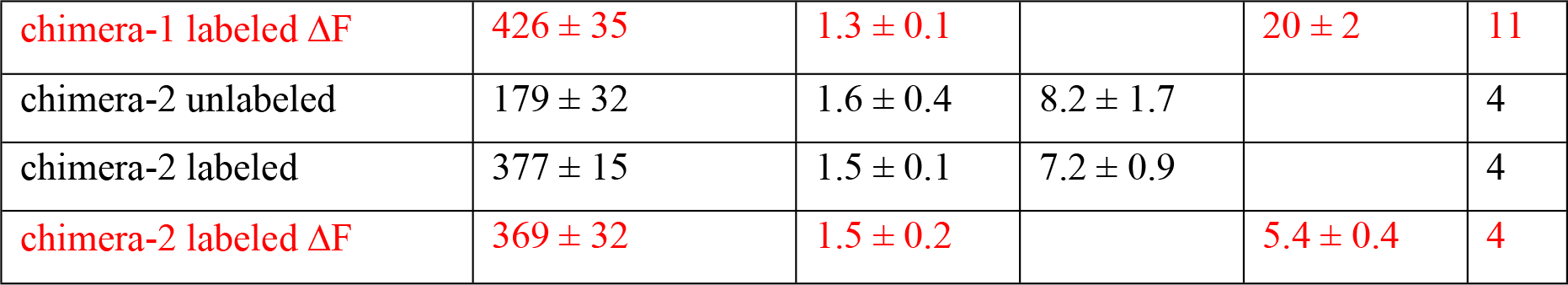

We have previously shown that when the α1-N203C or α3-N203C GlyRs are labelled by MTS-TAMRA, glycine application elicits large, reversible increases in ΔF [28, 34]. An example of a glycine ΔI and ΔF dose-response relationship recorded in an oocyte expressing the MTS-TAMRA-labelled α1-N203C GlyR is shown in Fig. 1E with the mean dose-response relationships plotted in Fig. 1G. The corresponding experiment for the MTS-TAMRA-labelled α2-N210C GlyR also yielded robust ΔI responses but no detectable ΔF (Fig. 1F, G). Mean parameters of best fit to all dose-response relationships are summarized in Table 1.

Similarly, we have previously shown that the α1-R271C and α3-R271C GlyRs are productively labelled by MTSR [28, 34]. Mean ΔI and ΔF dose-response relationships for the MTSR-labeled α1-R271C and α2-R278C GlyRs are shown in Fig. 1H with parameters of best fit summarized in Table 1. Again, whereas α1-R271C GlyRs yielded robust ΔF responses, α2-R278C GlyRs did not. Together, these results strongly suggest, despite their high amino acid sequence identities, significant structural differences exist in the N-terminal domains of α1-WT and α2-WT GlyRs.

To narrow down the receptor region responsible for the differential wF response, we generated a chimera (termed chimera-1) that comprised the N-terminal ligand-binding domain of the α1-N203C GlyR and the transmembrane domains and C-terminal tail of the α2-WT GlyR (Fig. 2A). A sample glycine gI dose-response of the unlabeled chimera-1 is shown in Fig. 2B. As seen in the mean dose-response curves presented in Fig. 2C, the glycine ΔI dose-response relationships for the α2-WT GlyR and chimera-1 were not significantly different. Following MTS-TAMRA labelling, the glycine ΔI EC_50_ was unchanged, suggesting minimal structural impairment, although the glycine ΔF dose-response now yielded a robust ΔF_max_ of 20 ± 2 % (n = 11, Table 1). However, we reasoned chimera-1 was too structurally distinct from the α2-WT GlyR to confidently infer the effects of phosphorylation on the α2-WT GlyR glycine binding site. Nevertheless the result indicated that the α2-WT GlyR N-terminus houses the domain responsible for the structural change that eliminates ΔF responses in MTS-TAMRA-labelled α2-N210C mutant GlyRs.

**Fig 2.**
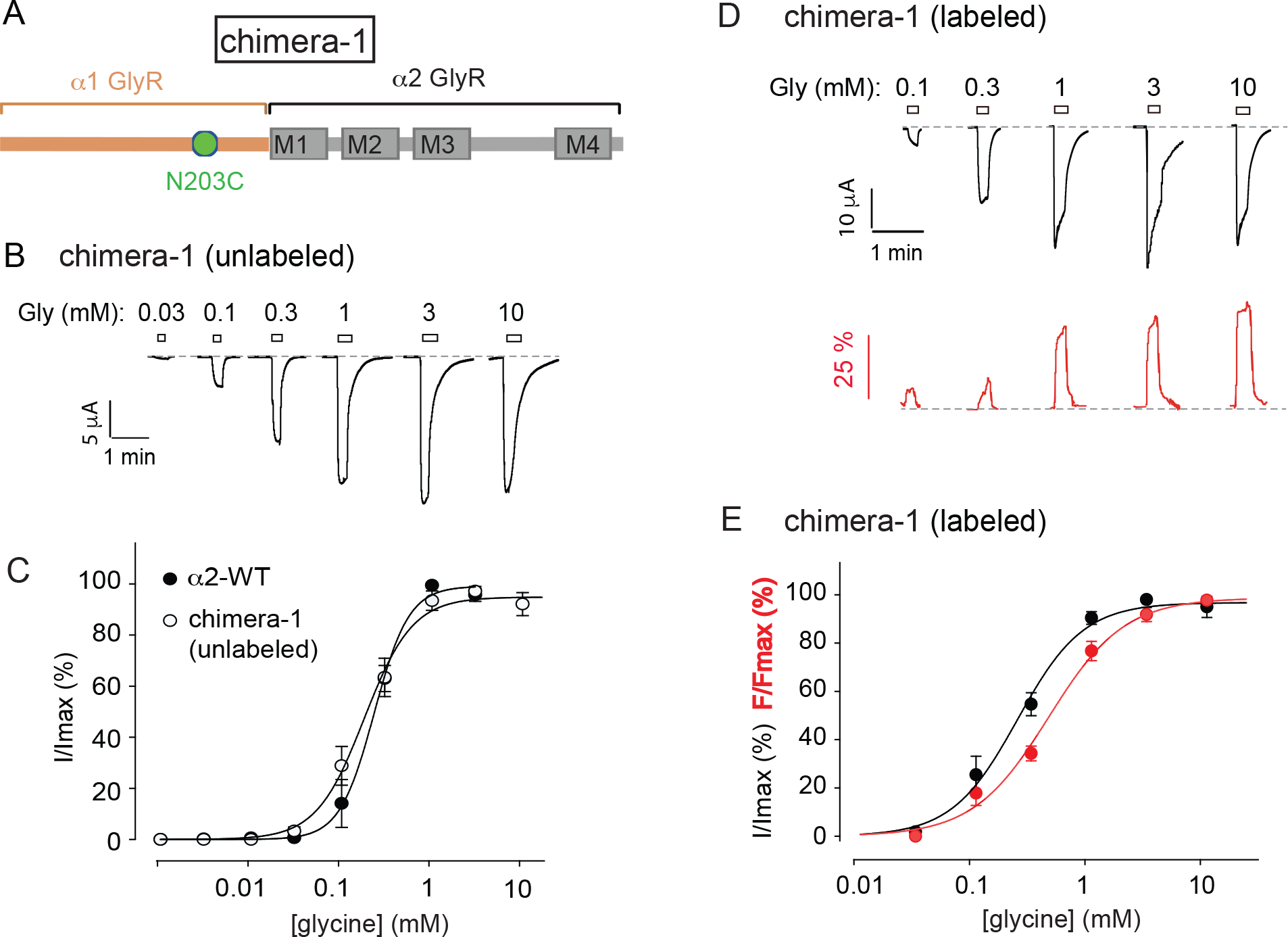
ΔI and ΔF activation properties of chimera-1 expressed in oocytes. **A.** Schematic model of chimera-1. Orange and gray coloring denote segments from the α1 and α2 GlyR sequences, respectively. **B.** Example of a chimera-1 glycine EI dose-response relationship. **C.** Averaged ΔI dose-response relations for the α2-WT GlyR and chimera-1. Mean parameters of best fit to individual dose-response relations are summarized in Table 1. **D.** Examples of glycine EI and ΔF dose-response relationships recorded from an oocyte that expressed MTS-TAMRA-labeled chimera-1. **E.** Averaged ΔI and ΔF dose-response relations for MTS-TAMRA-labeled chimera-1. Mean parameters of best fit to individual dose-response relations are summarized in Table 1.

The N-terminal domains of the α1-WT and α2-WT GlyRs are highly conserved. The region of lowest sequence identity is contained within binding domain loop F, also known as the β8-β9 loop. As part of the glycine binding site [35, 39], loop F is positioned at the extracellular subunit interface where it could interact directly with a label attached to binding domain loop C of the neighboring subunit (Fig. 1A). We thus hypothesized that non-conserved loop F residues were responsible for eliminating the sF response of α2-N210C GlyRs. To test this, we generated a new chimera, termed chimera-2, that comprised the α2-N210C GlyR with its loop F replaced by that of the α1-WT GlyR (Fig. 3A). This resulted in a total of five non-conserved residues between the α2-N210C GlyR and chimera-2 (Fig. 3A, inset). A sample glycine ΔI dose-response of the unlabeled chimera-2 is shown in Fig. 2B. As seen in the averaged data presented in Fig. 2C, the α2-WT GlyR and chimera-2 glycine ΔI dose-response relationships overlapped, implying minimal impact of the loop F structural changes on glycine binding interactions. Following MTS-TAMRA labelling the glycine ΔI dose-response of chimera-2 was unchanged, again implying minimal structural disruption, although now the ΔF dose-response yielded a ΔF_max_ of 5.4 ± 0.4 % (n = 4) (Fig. 3D, E; Table 1). As with the correspondingly labelled α1-N203C and α3-N203C GlyRs [28, 34], the MTS-TAMRA-labelled chimera-2 was also found to elicit a potent RF_max_ response (3.3 ± 0.7 %; n = 6) to a saturating (10 μM) concentration of strychnine. Together, these results suggest that MTS-TAMRA labelled chimera-2 may be useful for probing phosphorylation-induced structural changes in the α2-WT GlyR glycine binding site.

**Fig 3.**
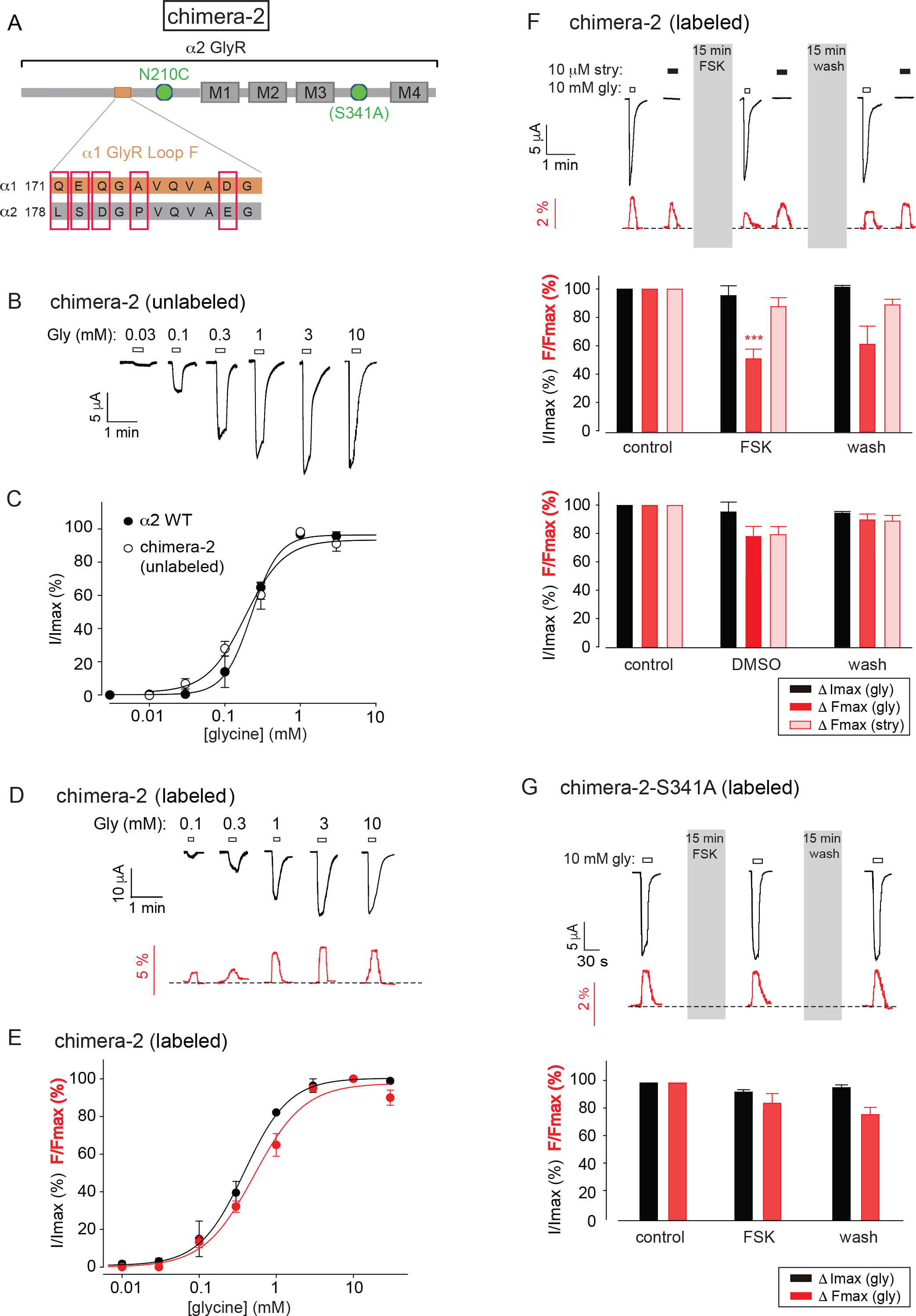
ΔI and ΔF activation properties of chimera-2 and the effect of phosphorylation when expressed in oocytes. **A.** Schematic model of chimera-2. Orange and gray coloring denote α1 and α2 GlyR segments, respectively. Two versions of chimera-2 were generated: with and without the S341A mutation. **B.** Example of a chimera-2 glycine ΔI dose-response relationship. **C.** Averaged ΔI dose-response relations for the α2-WT GlyR and chimera-2. Mean parameters of best fit to individual dose-response relations are summarized in Table 1. **D.** Examples of glycine ΔI and ΔF dose-response relationships recorded from an oocyte that expressed MTS-TAMRA-labeled chimera-2. **E.** Averaged ΔI and ΔF dose-response relations for MTS-TAMRA-labeled chimera-2. Mean parameters of best fit to individual dose-response relations are summarized in Table 1. **F.** The top panel shows examples of ΔI and ΔF responses mediated by MTS-TAMRA-labeled chimera-2 in response to saturating applications of glycine (10 mM) and strychnine (10 μM) before and after a 15 min forskolin (FSK) treatment and after a 15 min wash (n = 5 oocytes). The center panel shows averaged data for the experiment shown in the top panel (n = 5 oocytes), with all data points normalized to the respective control values recorded prior to FSK application. The bottom panel shows the effects of a control 15 min dimethyl sulfoxide (DMSO) treatment in place of FSK (n = 4 oocytes). *** p < 0.001 compared to control using paired t-test. **G.** Identical experiment to F except that the chimera-2 phosphorylation site has been eliminated by the S341A mutation (n = 6 oocytes).

### Phosphorylation of chimera-2 as detected by VCF

Based on the results of our previous study on the MTS-TAMRA-labelled α3-N203C GlyR [28], we hypothesized that phosphorylation of S341 may alter the magnitude of the glycine-induced ΔF_max_ of the MTS-TAMRA-labelled chimera-2. We tested this hypothesis directly by exposing oocytes expressing MTS-TAMRA-labelled chimera-2 to 20 μM forskolin for 15 min to in an attempt to phosphorylate S341. As shown in the sample recording in Fig. 3F (top and centre panels), it significantly reduced the glycine-mediated MF_max_ without affecting the strychnine-mediated ΔF_max_ (n = 5 oocytes). A control dimethyl sulfoxide application to the same labelled receptors had no effect on I_max_ or ΔF_max_ values (n = 4 oocytes), thus ruling out solvent-dependent effects (Fig. 3F, bottom panel). A second control experiment revealed that forskolin produced no significant change in ΔF_max_ magnitude in phosphorylation-deficient (S341A) mutant chimera-2 receptors (Fig. 3G), ruling out the possibility of non-specific forskolin effects on ΔF_max_ responses (n = 6 oocytes). As the glycine ΔF_max_ magnitudes remained constant throughout these experiments (Fig. 3F-G), we can also rule out an effect of phosphorylation on GlyR surface expression levels. Together, these results indicate that phosphorylation of S341 induced a conformational change in the immediate vicinity of the label attached to N203C in chimera-2.

### Phosphorylation of a2 GlyRs expressed in HEK293 cells

Given that S346 phosphorylation inhibits α3 GlyRs when expressed in mammalian HEK293 cells but not when expressed in *Xenopus* oocytes [28], it is evident that this effect can be expression system-specific. It was thus important to determine the effects of PKA-dependent phosphorylation on the functional properties of α2-WT GlyRs expressed in HEK293 cells. Note that it is not feasible to perform VCF on receptors expressed in HEK293 cells due to the high level of non-specific fluorophore labelling in HEK293 cell membranes. Our first experiment involved applying 20 μM forskolin for 15 min to α2-WT GlyRs that were subject to brief (2 s) applications of EC_100_ (3 mM) and EC_50_ (100 μM) glycine concentrations every 60 s. In a total of 16 cells we observed no significant effect of forskolin on current magnitudes activated by either EC_100_ or EC_50_ glycine concentrations. Although we supplemented the pipette solution with 2 mM Mg-ATP and 0.5 mM Na-GTP (see Methods) in these experiments, it remains possible that the lack of forskolin effect may have been due to insufficient levels of essential second messenger components in our HEK293 cell strain.

We thus adopted an alternate approach whereby we compared the functional properties of α2-WT GlyRs with those of phosphorylation-deficient α2-S341A GlyRs, and phosphorylation-mimicking α2-S341E GlyRs. Sample glycine dose-response experiments for the three receptors are shown in Fig. 4A. As shown in Fig. 4B and C and summarised in Table 2, the averaged glycine EC_50_ values, n_H_ values and I_max_ values were not significantly different from each other. Furthermore, when we applied a saturating (3 mM) glycine concentration for 25 s, the macroscopic decay rates of currents mediated by all three constructs were not significantly different from each other (Fig. 4D, E). However, we acknowledge that subtle changes in current activation and deactivation rates are not readily observable with standard whole-cell recording.

**Table 2.** Glycine-dependent activation properties of WT and mutant α2 GlyRs expressed in HEK293 cells

| Construct | EC<sub>50</sub> (μM) | $\eta_H$ | I<sub>max</sub> (nA) | decay time constant (s) | n |
| --- | --- | --- | --- | --- | --- |
| $\alpha 2$-WT | 153 ± 20 | 2.4 ± 0.2 | 5119 ± 501 | 4.85 ± 0.42 | 10 |
| $\alpha 2$-S341A | 142 ± 19 | 2.6 ± 0.2 | 6083 ± 1029 | 4.69 ± 0.46 | 10 |
| $\alpha 2$-S341E | 142 ± 15 | 2.4 ± 0.2 | 8543 ± 535 | 4.59 ± 0.48 | 12 |

**Fig 4.**
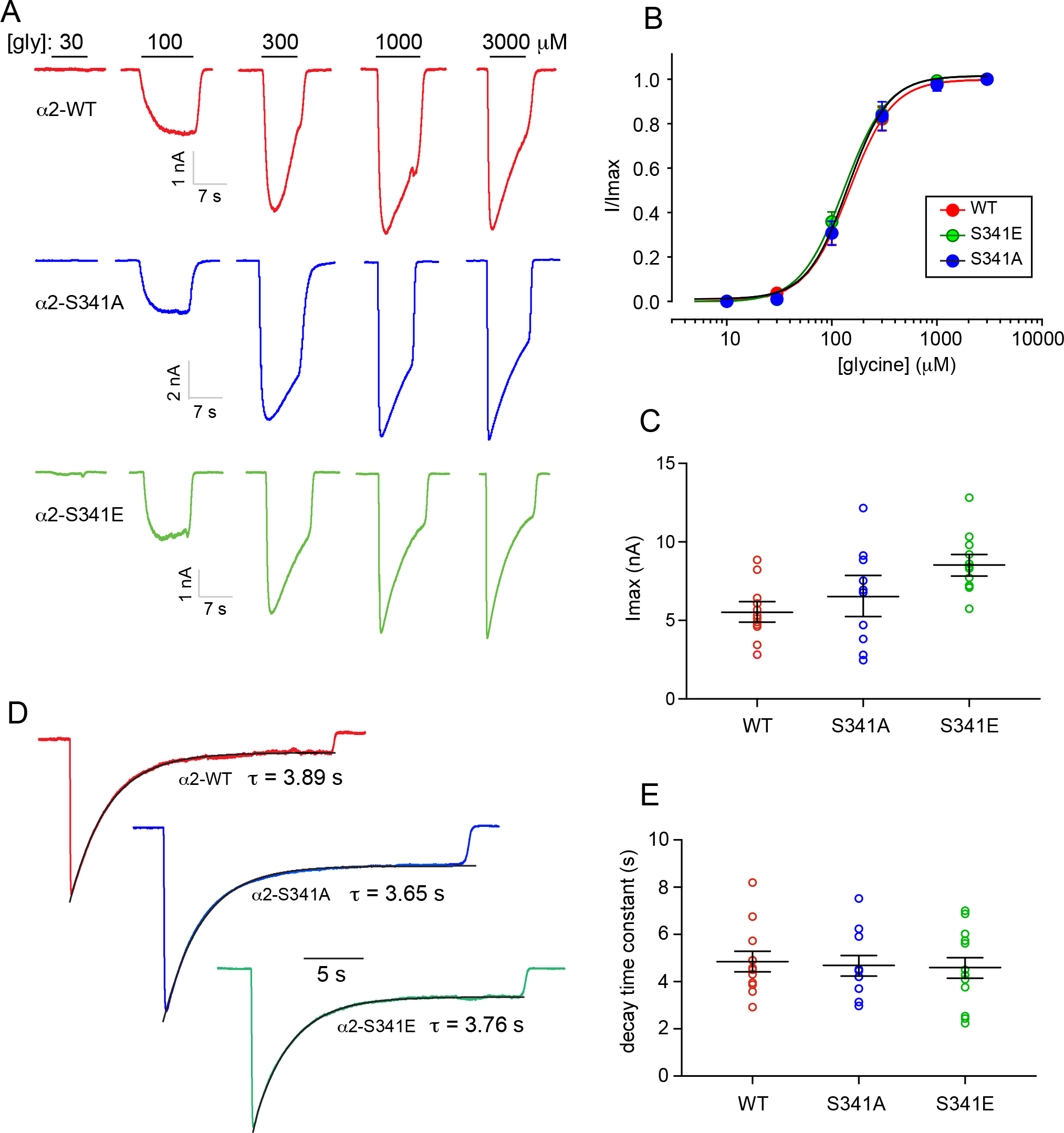
Functional analysis of α2-WT, α2-S341A and α2-S341E GlyRs in HEK293 cells. **A.** Glycine dose-response sample traces for the indicated GlyRs. **B.** Normalized, averaged glycine dose-response results for the indicated human GlyRs. Parameters of best fit to the Hill equation are summarized in Table 2. **C.** Mean I_max_ values for the three receptors. **D.** Comparison of macroscopic desensitisation rates of currents activated by saturating (3 mM) glycine. Mono-exponential curve fits to the current decay phases are indicated together with the fitted time constants. **E.** Mean decay time constants for the experiment shown in panel D.

Accordingly, we next investigated the properties of inhibitory postsynaptic currents (IPSCs) mediated by α2-WT, α2-S341A and α2-S341E GlyRs in artificial synapses. Whole-cell recordings from transfected HEK293 cells in co-culture with spinal neurons regularly exhibited robust, spontaneous IPSCs mediated by each of the three constructs (Fig. 5A). The advantage of this approach over transfected native neurons is that we can be sure that IPSCs are mediated by a pure population of α2 GlyRs. Fig. 5B shows digitally averaged and normalised IPSCs from single HEK293 cells expressing the WT and each mutant GlyR. Mean IPSC 10-90% rise times, amplitudes and decay time constants are presented in Figs. 5C-E. The respective parameters for the α2-WT GlyR correspond closely to those reported previously [31]. Although IPSC amplitudes did not vary significantly among the three receptors, the phosphorylation-mimicking α2-S341E GlyRs exhibited significantly slower IPSC rise and decay times relative to α2-WT and α2-S341A GlyRs (Fig. 5C, E).

**Fig 5.**
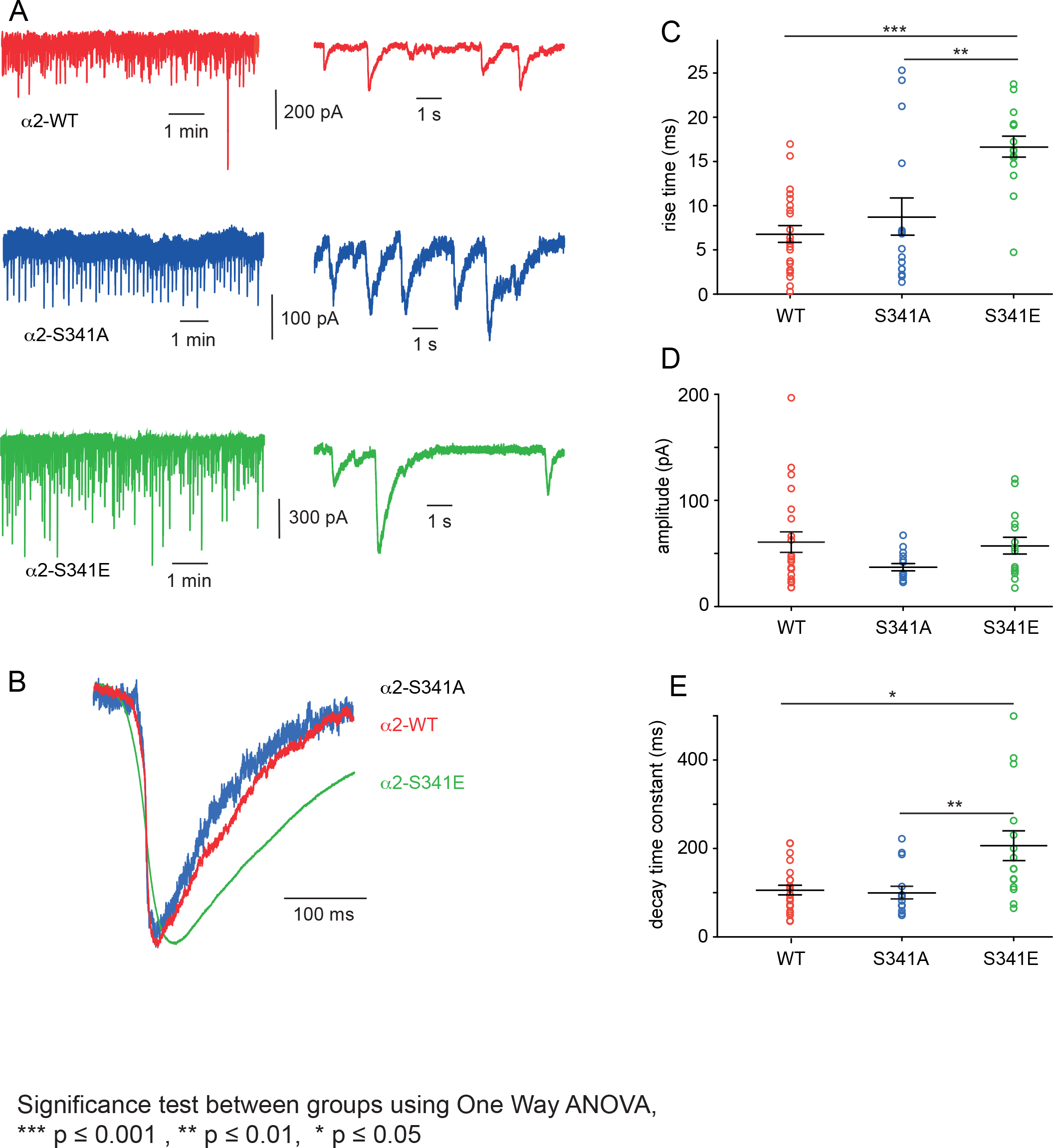
Properties of spontaneous IPSCs recorded from artificial synapses incorporating α2-WT, α2-S341A or α2-S341E GlyRs. **A.** Representative recordings of IPSCs from HEK293 cells expressing each isoform at two temporal scales. **B.** Averaged (from 50 - 100 events), normalized IPSCs from individual cells expressing each isoform. **C-E.** Mean IPSC 10 - 90% rise times, amplitudes and decay time constants. Significance was assessed using one way ANOVA, * p < 0.05, ** p < 0.01, *** p < 0.001.

## Discussion

We conclude that PKA-dependent phosphorylation of the α2 GlyR elicits two separable effects. The first effect, observed in artificial synapses, is a slowing in the IPSC rise and decay rates. As this was inferred from phospho-mimetic mutagenesis, it remains to be confirmed in native α2 GlyRs in neurons. The second effect of phosphorylation, observed in the *Xenopus* oocyte expression system, is a conformational change in or near the glycine-binding site. The main line of evidence in support of this is a forskolin-mediated change in the micro-environment of a fluorophore attached to loop C of the glycine-binding site, that is not observed when the phosphorylation site is eliminated. Note that phosphorylation modulated the glycine-induced ΔF_max_ but not the strychnine-induced ΔF_max_. Such a differential effect is not surprising given that the two ligands bind in different orientations and induce distinct conformational changes in the glycine binding site [33, 35].

We found that forskolin elicited no apparent effect on α2 GlyRs when recombinantly expressed in HEK293 cells. One possibility is that HEK293 cells did not have appropriate levels of expression of one or more intracellular signaling molecules essential for the phosphorylation process. Another possibility is that a phosphorylation-mediated binding site conformational change did occur but elicited no change in glycine sensitivity or current magnitude. Given that S341 may not have actually been phosphorylated by forskolin in our HEK293 cell experiments, we adopted the alternate approach of investigating the effects of phosphorylation-deficient (α2-S341A) and phosphorylation-mimicking (α2-S341E) mutations. Although we observed no effect of the mutations on glycine EC_50_, I_max_ or macroscopic decay rates in conventional whole-cell recordings (Fig. 4), we found that homomeric α2-S341E GlyRs mediated IPSCs with significantly slower rise and decay times in artificial synapses (Fig. 5).

The α2 GlyR is initially expressed as a homomer in embryonic neurons [40] and its expression in these neurons has long been known to be important for synaptogenesis [41–43], cortical development [44] and retinal photoreceptor development [45]. Because the expression of the potassium-chloride cotransporter type-2 (KCC2) is low in embryonic neurons [46], the intraceullar chloride concentration is high and as a result, α2 GlyR activation is excitatory [47]. In embryonic migrating cortical interneurons, activation of α2 GlyRs thus leads to neuronal depolarisation, the subsequent activation of a voltage-gated calcium influx and to the modulation of spontaneous calcium oscillations [19]. These in turn modulate myosin activity that alters interneuron nucleokinesis and increases interneuron migration speed [19]. Functional α2 GlyRs also contribute to the differentiation of cortical neuroprogenitors into functioning neurons[20]. Thus, α2 GlyRs are essential for the development of correctly functioning cortical circuits. Conversely, loss or dysfunction of α2 GlyRs is associated with neurodevelopmental disorders including schizophrenia, autism and epilepsy [25] [26, [50] [51].

Although it undergoes a decline in expression levels during development[40], the α2 GlyR remains widely distributed throughout the adult central nervous system [40, 52]. Indeed it has recently been shown that homomeric α2 and α3 GlyRs mediate tonic glycine-mediated currents throughout higher brain regions [21] [53]. Homomeric GlyRs are also found presynaptically at cortical glutamatergic synapses [51, 54-56]. These presynaptic GlyRs are likely to be both α2 and α3 GlyRs, given that 1) the α2 GlyR has been positively identified in these locations [51] [55] and 2) a missense mutation (α2-P192L, α2-P185L) common to both subunits results in a common temporal lobe epilepsy phenotype [50]. When expressed presynaptically, homomeric α2 GlyRs would be exposed to phasically released glycine. In this case, PKA phosphorylation would prolong receptor open times and thereby prolong the depolarization of presynaptic terminals. This is likely to increase calcium influx, thereby enhancing neurotransmitter release and leading to enhanced excitability of postsynaptic neurons.

In summary, we have demonstrated that phosphorylation of S341 exerts a conformational change that propagates to the α2 GlyR glycine-binding site. We also found that the S341E phospho-mimetic mutation results in slower IPSC rise and decay times. These results suggest that PKA phosphorylation can alter the structure and functional properties of the α2 GlyR. This information adds to the complexity of glycinergic modulatory mechanisms, and may help to identify new mechanisms by which α2 GlyRs may be pathologically modified or therapeutically targeted for the treatment of neurological disorders including epilepsy.

## Acknowledgements

This study was supported by a grant from the National Health and Medical Research Council of Australia (1058542) to JWL.

**Author contributions**
S.I. – performed experiments, analysed and interpreted data
X.C. – performed experiments, analysed and interpreted data
A.E.-M. – performed molecular modelling
J.W.L. – designed project, performed experiments, interpreted data, wrote manuscript

